# Lightway access to AlphaMissense data that demonstrates a balanced performance of this missense mutation predictor

**DOI:** 10.1101/2023.10.30.564807

**Authors:** H. Tordai, O. Torres, M. Csepi, R. Padányi, G. L. Lukács, T. Hegedűs

## Abstract

Single amino acid substitutions can profoundly affect protein folding, dynamics, and function, leading to potential pathological consequences. The ability to discern between benign and pathogenic substitutions is pivotal for therapeutic interventions and research directions. Given the limitations in experimental examination of these variants, AlphaMissense has emerged as a promising predictor of the pathogenicity of single nucleotide polymorphism variants. In our study, we assessed the efficacy of AlphaMissense across several protein groups, such as mitochondrial, housekeeping, transmembrane proteins, and specific proteins like CFTR, using ClinVar data for validation. Our comprehensive evaluation showed that AlphaMissense delivers outstanding performance, with MCC scores predominantly between 0.6 and 0.74. We observed low performance on the CFTR and disordered, membrane-interacting MemMoRF datasets. However, an enhanced performance with CFTR was shown when benchmarked against the CFTR2 database. Our results also emphasize that quality of AlphaFold’s predictions can seriously influence AlphaMissense predictions. Most importantly, AlphaMissense’s consistent capability in predicting pathogenicity across diverse protein groups, spanning both transmembrane and soluble domains was found. Moreover, the prediction of likely-pathogenic labels for IBS and CFTR coupling helix residues emphasizes AlphaMissense’s potential as a tool for pinpointing functionally significant sites. Additionally, to make AlphaMissense predictions more accessible, we have introduced a user-friendly web resource (https://alphamissense.hegelab.org) to enhance the utility of this valuable tool. Our insights into AlphaMissense’s capability, along with this online resource, underscore its potential to significantly aid both research and clinical applications.

## Introduction

In both the medical field and the broader realm of biology, understanding the pathogenicity of mutations holds high significance (1,2). Pathogenic mutations disrupt the normal function of genes, leading to multiple diseases and medical conditions. From the early onset of genetic disorders in infants to the development of complex diseases in adults, the transformative power of a single nucleotide change can be profound. Discerning between benign and pathogenic mutations can influence diagnostic accuracy, guide therapeutic interventions, and inform prognosis (3). Therefore, reliable tools and methodologies to predict and understand mutation impact are essential.

Prior to the advent of more advanced genetic analytical tools, several algorithms emerged as standard bearers in predicting the potential impact of mutations, such as PROVEAN, PolyPhen-2, and SIFT. PROVEAN (Protein Variation Effect Analyzer) offers predictions based on the alignment of homologous protein sequences. Meanwhile, PolyPhen-2 (Polymorphism Phenotyping v2) employs a combination of sequence and structural information to classify variants as benign or probably damaging (4). SIFT (Sorting Intolerant From Tolerant) operates by considering the degree of conservation of amino acid residues in sequence alignments derived from closely related sequences to predict whether an amino acid substitution affects protein function (5). While these tools have undeniably advanced our understanding of mutation pathogenicity, they also underscore the complexity of the task and highlight the need for continuous refinement in the face of rapidly accumulating genomic data. Newer tools for evaluating the pathogenicity of missense mutations evolved. MVP (Missense Variant Pathogenicity prediction) has gained attention for its sophisticated integration of multiple features related to genetic variation (6). MetaSVM is an ensemble method that merges the outputs of various tools using support vector machines to consolidate pathogenicity prediction (7). M-CAP (Mendelian Clinically Applicable Pathogenicity) stands out for its high specificity in distinguishing disease-associated variants from neutral ones (8). VESPA, the Variant Effect Scoring Prediction Algorithm, is based on embeddings of a protein language model, which captures nuanced relationships between amino acid residues, allowing for a more refined and context-aware prediction of variant impacts (9).

AlphaMissense machine learning, developed recently by DeepMind, can predict the pathogenicity of missense variants and stands at the frontier of missense variant pathogenicity prediction (10). Importantly, it leverages the structural prediction capabilities of AlphaFold (11) to analyze these variants. To potentially enhance the precision of missense variant pathogenicity insights, AlphaMissense evolved the field by merging sophisticated machine learning with structural biology. Moreover, AlphaMissense aims to tackle the challenge of interpreting the vast number of missense variants in the human genome, many of which have unclear clinical significance. It holds the promise of revolutionizing the understanding and diagnosis of genetic diseases by classifying missense variants as likely benign or likely pathogenic (10).

While the conception of AlphaMissense represents a commendable stride, defined by its intricate design and advanced methodologies, there remain gaps in our understanding of its performance on selected groups of proteins or individual proteins. In particular, a pivotal concern arises from the specificities of its missense mutation predictions and the limited accessibility to its dataset. Whereas there are initiatives to make the data accessible through R and Python tools (12–16), these require a certain level of computational skills, thus significantly restricting the user base. Addressing these voids, we assessed AlphaMissense performance on different datasets using ClinVar data and developed a web resource that notably facilitates streamlined data extraction for research purposes and also offers a visual representation of these predictions within a structural context.

## Methods

### Datasets

The primary AlphaMissence dataset, AlphaMissense_hg38.tsv.gz, was sourced from Zenodo (https://zenodo.org/records/8208688). This data contains all predictions with all possible missense variations in the human proteome. Missense data was retrieved from ClinVar (17) as of 26^th^ September 2023 and made available at Zenodo (https://doi.org/10.5281/zenodo.10023060). The dataset representing the human proteome was obtained from UniProt Release 2023_04, specifically from the file UP000005640_9606.dat (reference proteomes from https://www.uniprot.org/help/downloads) (18). This dataset proved instrumental in mapping Ensemble IDs from ClinVar to UniProt accession numbers since the inherent online ID mapping tool at UniProt matched only a very low number of entries. Human protein Structures were downloaded from AlphaFoldDB (version 4; https://alphafold.ebi.ac.uk/download#proteomes-section) (19). Mitochondrial Protein Data was procured from MitoCharta (https://www.broadinstitute.org/mitocarta/mitocarta30-inventory-mammalian-mitochondrial-proteins-and-pathways) (20). The boundaries of membrane regions in transmembrane proteins were sourced from the Human Transmembrane Proteome (HTP; https://htp.unitmp.org) and filtered to include only entries boasting a quality score greater than 85 to maintain the integrity and accuracy of our analyses (21). Entries omitted from the TM analysis were not incorporated into the dataset encompassing soluble or membrane-associated proteins. However, this criterion resulted in a sparse representation of high-quality predictions for ABC proteins. Therefore, we supplemented the data with TM boundaries from our proprietary ABCM2 database (http://abcm2.hegelab.org) (22,23).

### Analysis

All data analyses were carried out using Python-based tools to ensure flexibility and scalability. The core dataset of AlphaMissense is housed within a Postgresql database (https://www.postgresql.org), a robust and efficient system suitable for handling extensive datasets. To facilitate a lightweight and seamless interaction with the stored data, we employed the SQLalchemy library (24) renowned for its capability to provide a high-level, Pythonic interface to relational databases.

Matplotlib was used for generating plots that delineate various aspects of the data (25). Structural visualization of proteins was done using PyMOL (version 2.4, Schrödinger, LLC.), a molecular graphics system with an embedded Python interpreter. To bridge the predictions of AlphaMissense with these structures, MDAnalysis was employed (26). This Python toolkit allowed us to incorporate the AlphaMissense scores directly into the PDB files, specifically inserting them into both the occupancy and B-factor columns.

### Web application

We incorporated robust technologies tailored for optimal performance and interactivity to deliver a seamless, responsive, and user-friendly experience for users accessing our web resource. At the heart of our web application lies FastAPI (https://fastapi.tiangolo.com), a high-performance web framework for building APIs with Python. For rendering dynamic content on the front end, we employed Jinja2, offering the flexibility to inject real-time data into our HTML templates, facilitating a dynamic user interface. The visual presentation and interactivity of the web resource are bolstered by Bootstrap 5 to bring enhanced responsiveness, aesthetic components, and functionality. In sync with our data analysis stack, the web resource continues to leverage Postgresql for data storage. To render intricate molecular structures directly within the browser, we integrated PDBe-molstar (27). This state-of-the-art WebGL component ensures high-quality, interactive 3D visualizations, allowing users to engage deeply with the structural context of the AlphaMissense predictions.

## Results and Discussion

### Performance of AlphaMissense across diverse protein groups in relation to ClinVar data

The performance of AlphaMissense may exhibit variability across different protein types, necessitating careful scrutiny when analyzing target proteins. We evaluated AlphaMissense’s efficiency across a range of protein groups, choosing ClinVar as our benchmark. While ClinVar is a valuable resource, it has its shortcomings. For instance, it may disproportionately represent genes under intensive study while underrepresenting highly pathogenic mutations due to the fact that individuals harboring them might not survive to birth. For our analysis, we juxtaposed all benign and pathogenic missense mutations rated with at least one star in ClinVar against AlphaMissense predictions for proteins in our datasets. Only genes with corresponding ClinVar entries were considered. Subsequently, we derived precision, recall, F1 score, aucROC, and Matthew’s Correlation Coefficient (MCC) (Table 1). In general, the calculated statistical measures were high for all of the groups studied. Most importantly, MCC exceeded 0.6 for all but two groups, with low values possibly stemming from sparse input data for MemMoRFs and compromised ClinVar data quality, especially for CFTR.

**Table 1:** AlphaMissense performance on proteins sets, benchmarked with ClinVar.

| | $n(\text{protein})^1$ | $n(\text{mutation})^2$ | $PPV^4$ | $TPR$ | $F1$ | $aucROC$ | $MCC$ | $f(CV, \text{benign})^5$ | $f(CV, \text{pathogenic})$ | $f(AM, \text{benign})$ | $f(AM, \text{pathogenic})$ |
| --- | --- | --- | --- | --- | --- | --- | --- | --- | --- | --- | --- |
| *ALL*<sup>3</sup> | 11,486 | 107,681 | 0.829 | 0.776 | 0.802 | 0.870 | 0.697 | 0.009 | 0.005 | 3.425 | 1.713 |
| *MITO* | 299 | 2,970 | 0.804 | 0.874 | 0.837 | 0.888 | 0.702 | 0.011 | 0.010 | 3.085 | 2.039 |
| *HK* | 1,011 | 8,329 | 0.878 | 0.801 | 0.838 | 0.861 | 0.714 | 0.008 | 0.006 | 2.899 | 2.285 |
| *SOL* | 7,938 | 73,816 | 0.831 | 0.746 | 0.786 | 0.867 | 0.686 | 0.009 | 0.004 | 3.484 | 1.755 |
| *IBS* | 138 | 55 | 0.895 | 0.944 | 0.919 | 0.890 | 0.755 | 0.340 | 0.760 | 1.800 | 3.880 |
| *MemMoRF* | 35 | 21 | 0.462 | 1.000 | 0.632 | 0.846 | 0.496 | 0.381 | 0.619 | 2.286 | 2.619 |
| *Low-AF* | 12 | 327 | 0.828 | 0.358 | 0.500 | 0.831 | 0.481 | 0.022 | 0.002 | 2.466 | 1.952 |
| *HTP85* | 1,653 | 16,015 | 0.815 | 0.855 | 0.834 | 0.872 | 0.704 | 0.009 | 0.007 | 3.352 | 1.762 |
| *HTP85 – TM* | 1,653 | 1,997 | 0.895 | 0.900 | 0.898 | 0.872 | 0.745 | 0.453 | 0.683 | 2.127 | 3.045 |
| *HTP85 – nonTM* | 1,653 | 14,018 | 0.799 | 0.845 | 0.822 | 0.870 | 0.694 | 0.623 | 0.471 | 3.336 | 1.881 |
| *GPCR* | 299 | 1,696 | 0.802 | 0.651 | 0.719 | 0.871 | 0.631 | 0.009 | 0.003 | 3.573 | 1.647 |
| *ABC* | 42 | 1,557 | 0.770 | 0.948 | 0.850 | 0.882 | 0.646 | 0.009 | 0.019 | 3.370 | 1.787 |
| *CFTR* | 1 | 207 | 0.884 | 1.000 | 0.939 | 0.975 | 0.478 | 0.005 | 0.134 | 3.289 | 1.776 |
| *CFTR (CFTR2)* | 1 | 119 | 0.961 | 0.961 | 0.961 | 0.852 | 0.725 | 0.011 | 0.069 | 3.289 | 1.776 |
| *lowAF* | 12 | 327 | 0.828 | 0.358 | 0.500 | 0.831 | 0.481 | 0.022 | 0.002 | 2.466 | 1.952 |
| *lowAF-pLDDT50* | 12 | 126 | 0.828 | 0.571 | 0.676 | 0.819 | 0.573 | 0.010 | 0.003 | 1.843 | 2.595 |
| *SOL-pLDDT50* | 7,938 | 44,045 | 0.832 | 0.785 | 0.808 | 0.843 | 0.660 | 0.007 | 0.005 | 2.825 | 2.333 |

<sup>1</sup> Number of proteins with associated ClinVar entries.
<sup>2</sup> Number of ClinVar mutations with at least one star, for the given set of proteins.
<sup>3</sup> MITO: mitochondrial, HK: housekeeping, SOL: soluble, IBS: interfacial binding site, HTP85: Proteins in the Human Transmembrane Proteome with at least a confidence score of 85, TM: transmembrane region only, lowAF: low quality AlphaFold structures, -pLDDT50: sets without residues with a pLDDT score lower than 50.
<sup>4</sup> PPV: positive predictive value, TPR: true positive rate, aucROC: Area Under the Receiver Operating Characteristic Curve, MCC: Matthews's correlation coefficient.
<sup>5</sup> The number of benign and pathogenic mutations from ClinVar (CV) data and from AlphaMissense (AM) predictions was normalized to the number of amino acids (summed length of proteins) for each protein set.

We also determined the frequency of likely benign and pathogenic mutations in ClinVar relative to protein length (Table 1). Our initial analysis centered on mitochondrial proteins of bacterial origin. Given the unique sequence attributes of these proteins, prediction biases were anticipated. Intriguingly, the pathogenic variation frequency for these proteins was higher than that of the entire human protein ensemble. The important cellular function of these proteins in energy balance might hint their role as housekeeping genes. Drawing from a specific database (https://housekeeping.unicamp.br) (28), we cross-referenced 1,011 housekeeping genes with 299 mitochondrial genes from our collection and only a modest overlap of 98 genes was observed. Interestingly, the anticipated elevation in pathogenic mutation frequency was as evident in the housekeeping gene dataset as expected.

We also investigated membrane-interacting residues. One dataset included interfacial binding site (IBS) residues (29) while the other contained membrane molecular recognition features (MemMoRFs; lipid-interacting disordered regions) (30). For IBS residues, pathogenic mutations were approximately twice as frequent as benign ones (0.760 vs. 0.340), likely reflecting the functional significance of these residues. Similar trends were evident for the MemMoRF set, although it’s crucial to recognize the limited sample size for this category that might explain the diminished MCC when comparing ClinVar and AlphaMissense outcomes. Moreover, the intrinsic disorder and low sequence conservation of these regions might also influence AlphaMissense’s predictive power on these proteins (10).

Mutation frequencies and AlphaMissense efficiency on transmembrane (TM) proteins were also assessed. We segregated residues into TM and non-TM subsets using the Human Transmembrane Proteome database (21). Counterintuitively, AlphaMissense performed better on TM regions, despite their hydrophobicity potentially reducing evolutionary insights from sequence alignments. However, the spatial constraints of transmembrane domains lacking intrinsically disordered regions might boost the AlphaFold-based AlphaMissense predictions (31). Remarkably, pathogenic mutations were more prevalent in TM domains than benign ones (Table 1).

We also focused on specific membrane protein subsets. While a surge in pathogenic mutations for GPCRs in ClinVar was anticipated, this was not observed. In contrast, ABC proteins manifested elevated pathogenic mutation frequencies in the ClinVar database. Such disparities might be the result of the disease linkage of specific protein classes or research biases. Importantly, type and quality of data can profoundly impact these types of analyses. For instance, when juxtaposing AlphaMissense’s predictions against ClinVar data for the CFTR/ABCC7 protein, benign mutations were infrequent, whereas pathogenic mutations predominated. The MCC for CFTR ClinVar/AlphaMissesnse comparison was low (0.478).

Finally, the potential source of low MCC values were investigated. In the case of CFTR, we tested AlphaMissense predictions against a gold standard CFTR mutation database, CFTR2 (The Clinical and Functional TRanslation of CFTR (CFTR2); available at http://cftr2.org). The CFTR2 database exhibited benign mutation frequencies comparable to other groups but a marked increase in pathogenic mutations. The calculated MCC with this benchmark set was one of the highest (0.725) compared to any of the other protein groups. We assumed that the very low MCC for MemMoRF groups may have caused by the high prevalence of disordered residues in these proteins. Because of the small size of this dataset we tested this possibility on soluble proteins, by excluding those residues from the calculations, which residues exhibit a pLDDT score lower than 50 in AlphaFold structures as a proxy for intrinsically disordered regions (32). A small increase was observed for PPV, TPR, and F1, but not for rocAUC and MCC values (Table 1). Therefore, we assumed that low results of proteins with MemMoRF may have arisen from the AlphaFold’s capabilities for predicting their structures, since they involve several single-pass, bitopic transmembrane proteins. We also indirectly investigated this possibility, and used a transmembrane protein set with failed AlphaFold predictions (33), which group of proteins (low-AF, Table 1) resulted also very low MCC scores. Interestingly, excluding residues with a pLDDT score lower than 50 increased the TPR, F1, and MCC scores. The latter score for this set became 0.573.

### Variability in AlphaMissense predictions across different groups of proteins

The observed differences in True Positive Rate (TPR) and F1 scores implied that the distribution of benign and pathogenic mutations is not uniform across protein groups. To gain a deeper insight and understand AlphaMissense’s predictive properties, we investigated the frequency and distribution of its predictions across various protein categories (Table 1). Typically, benign mutations were more frequent, with values hovering between 3 to 3.5, as opposed to pathogenic mutations, which ranged from approx. 1.5 to 1.8. However, certain intriguing trends emerged. For instance, housekeeping genes showcased slightly reduced benign and elevated pathogenic mutation frequencies. Mitochondrial proteins leaned towards a higher pathogenic frequency. A distinct trend was observed for the IBS dataset encompassing only functional sites containing IBS, their benign mutation frequency was highly decreased, while pathogenic mutation frequency was increased. A similar pattern was evident for transmembrane regions of transmembrane proteins. Given that AlphaMissense predictions cover all possible missense mutations, not biased by human issues, it is reasonable to deduce that only about 30-35% of the possible human missense mutations are pathogenic. This general trend was also mirrored in the mutation frequencies observed in the ClinVar data for certain protein categories (Table 1).

We next examined whether the reverse mutations demonstrated similar average AlphaMissense scores. For each variation, we calculated the mean scores and paired them with their reverse counterpart for visualization. We highlighted variation pairs that showed a difference of at least 0.2 in their average scores (Fig. 1a). The pathogenicity labels of three pairs are changed from pathogenic to benign (highlighted by asterisks). The contrasting mean values of the Cys/Ser mutation, categorized as likely-pathogenic, and the Ser/Cys, which is deemed likely-benign, can be rationalized based on amino acid properties and structural implications. Cysteine plays a pivotal structural role, particularly in forming disulfide bridges. In a simplified form, this makes the replacement of Serine with Cysteine more feasible than the other way around, as Serine cannot replicate Cysteine’s capability in forming disulfide bridges. Accordingly, Cys/Ser pathogenic mutation frequency (0.011) is 5.5 times higher than Ser/Cys pathogenic labels (0.002) in the ClinVar dataset.

**Figure 1:**
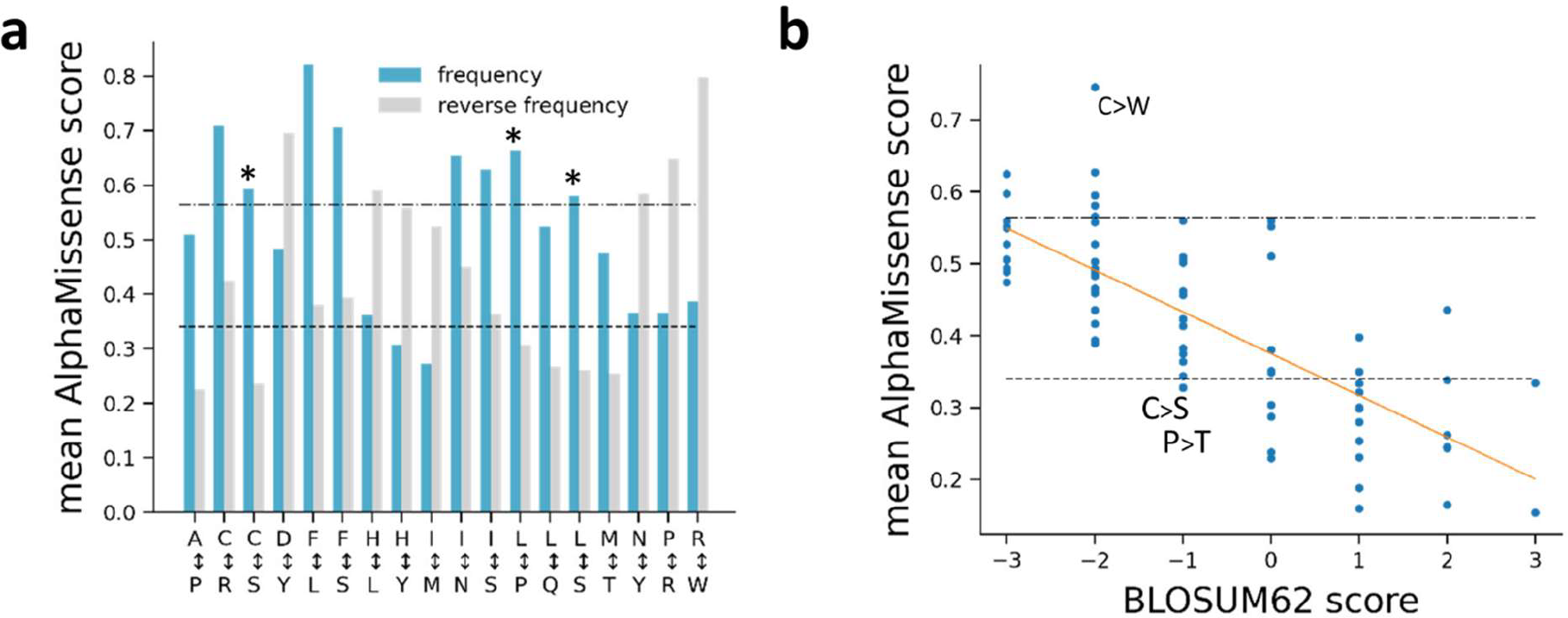
(**a**) Mean AlphaMissense scores for variations, which display a minimum score difference of 0.2 when compared to the reverse amino acid change. Asterisks mark those changes which get the opposite label (benign/pathogenic) in the case of reverse change. (**b**) Mean AlphaMissence scores for each variation grouped by their BLOSUM62 score. Dashed and dashed dotted lines indicate the cutoffs of the ambiguous AlphaMissense predictions. Orange line was fitted (r=-0.755, p=5.13×10^−15^).

We also analyzed how the mean scores of all variations correlated with the symmetric BLOSUM62 matrix, a representation derived from amino acid substitution frequencies based on sequence alignments. A noticeable correlation emerged: variations with lower BLOSUM62 scores tended to have higher AlphaMissense scores (correlation coefficient: -0.755, p=5.13×10^−15^, Fig. 2b). Most of the average scores for less favorable substitutions fell below the likely-pathogenic threshold set by AlphaMissense. This trend may arise from the higher ratio of variations predicted as likely-benign. Notably, the averages for both Cys/Ser and Ser/Cys variations, which have a BLOSUM62 substitution score of -1, lie slightly below 0.34, placing them in the likely-benign category. This observation also applies to the Pro/Thr pair (Fig. 2b). In contrast, the Cys/Trp pair, with a substitution score of -2, recorded the highest AlphaMissense score of 0.746. Substitutions with a score of -4 (Cys/Glu, Trp/Pro, Trp/Asn, Trp/Asp, and Phe/Pro) are not present, since they cannot be rationalized with a single nucleotide change.

**Figure 2:**
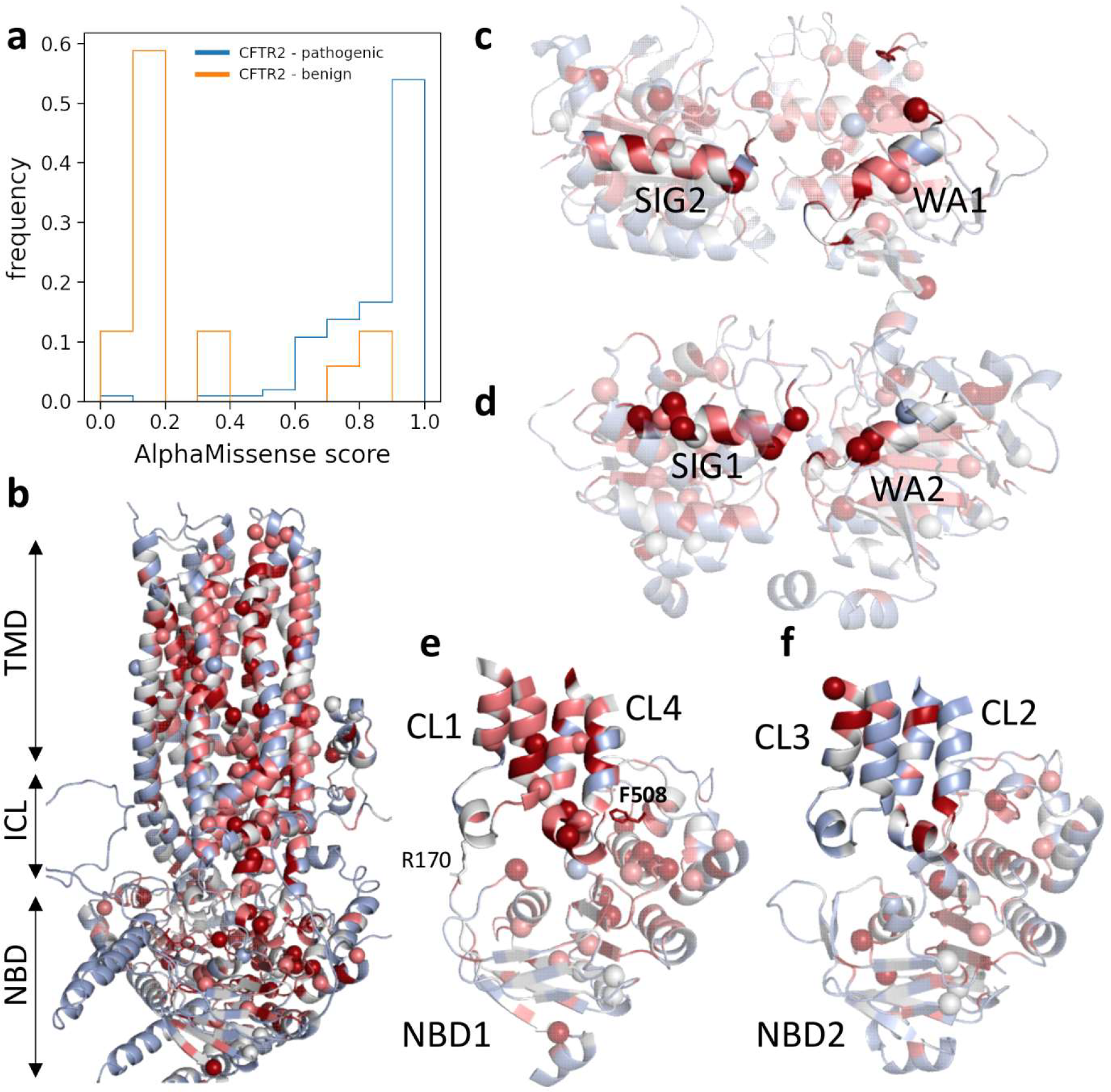
Distribution of AlphaMissense scores for CFTR. (**a**) Histograms of AlphaMissense scores for benign (n=20) and pathogenic (n=102) mutations from the CFTR2 database. (**b**) AlphaFold structure of CFTR colored by mean AlphaMissense scores. Blue: 0-0.340, gray: 0.340-0.564, pink: 0.564-0.780, red: 0.78-1. Spheres represent pathogenic mutations from CFTR2. (**c, d**) The degenerate, non-catalytic ATP-binding Site-1 and catalytic Site-2. Residues 461-472 and 1346-1362 were highlighted for Site-1 structural elements and residues 548-564 and 1247-1258 for Site-2. (**e, f**) Pathogenic mutations and mean scores at the NBD/TMD interfaces. TMD: transmembrane domains, ICL: intracellular loops, NBD: nucleotide binding domains, CL: cytoplasmic loops, sticks: F508 and R170. Coloring scheme of all structures is the same.

### Exploring the AlphaMissense prediction of the ABC transmembrane protein, CFTR

We assessed the AlphaMissense predictions for the CFTR protein, which attracted substantial attention within the scientific community, primarily because of its association with cystic fibrosis (34). For our study, we relied on the CFTR2 database to annotate mutations. Impressively, out of the 102 pathogenic and 20 benign mutations listed in the CFTR2 database, AlphaMissense mispredicted only four pathogenic (I601F, A613T, I1234V, and V1240G with scores 0.49, 0.39, 0.08, and 0.5637) and four benign (F508C, L997F, T1053I, and R1162L with scores 0.87, 0.74, 0.35, and 0.89) mutations to the opposite or ambiguous category. Performance metrics for AlphaMissence on CFTR are listed in Table 1 and the particular scores of the 102 values are visualized in Fig. 2a.

A spatial representation of these mutations on the CFTR structure reveals some intriguing patterns (Fig. 2b). The ATP binding sites of CFTR, especially, warrant attention. The formation of an ATP binding site is an intricate interplay between one Walker A motif from a Nucleotide Binding Domain (NBD) and a signature motif from the opposite NBD. In comparison to the functional site-2, both the count of CFTR2-sourced mutations and the AlphaMissense scores were observed to be lesser at the site1 (0.584 versus 0.493 and 15 versus 3, respectively; Fig. 2c,d), which is degenerate, rendering it incapable of ATP hydrolysis (35). Moreover, the structural landscape around the F508 residue provides more insight. The CL4 coupling helix, which interfaces with the F508 residue, presents a greater number of both predicted and CFTR2-based mutations in comparison to CL2 (as visualized in Fig. 2e,f), which is a structural counterpart of CL4. No CFTR2 mutations are present in other than CL4 coupling helix, CH4. CH1, 2, 3, and 4 mean AlphaMissense scores are 0.336, 0.411, 0.136, and 0.648, respectively. Interestingly, CL1 was found to be devoid of CFTR2 mutations, but in vitro experiments in this region revealed that the R170G mutation, which has a likely-benign AlphaMissense label, impairs the domain-domain assembly and would be pathogenic if harbored by an individual (36).

The F508 residue is not only an epicenter for deleterious mutations but has also been extensively researched. While CFTR2 lists no additional pathogenic mutations for this residue, a range of experimental works have delved into substituting the F with all the other 19 possible amino acids to discern the impacts on the functional expression of CFTR (37). Some of these substitutions, including F508I, F508L, F508V, F508C, F508S, and F508Y, are featured in the AlphaMissense dataset. Uniformly, they were all predicted as likely pathogenic. Nevertheless, apart from the F508C variant, experimental data suggests that the F508V mutation might also be functionally permissive (37), deviating from AlphaMissense’s likely-pathogenic prediction. Two other variants, labeled as “unknown” or of “varying significance” in the CFTR2 database, show discrepancies between *in vitro* experiments and AlphaMissense predictions. Specifically, the F1052V mutation, predicted by AlphaMissense as likely-pathogenic, demonstrates a functional expression, with 57% mature protein form and 60% functionality relative to the wild type (38). Conversely, the S912L variant, predicted as benign, appears to be a potential false negative AM prediction. This is based on its substantially reduced function, at 16% of the wild type, despite an expression level nearly on par at 92% relative to the wild type (38).

### A web resource to ease access and mutational hot spot detection

Our web resource was developed to offer users a simple and intuitive interface to delve deep into the AlphaMissense data. On the search page, the users access the flexibility of using a wide range of identifiers to query proteins. Whether it is UniProt ACC, Entry name, Gene name, or Ensembl transcript ID, the system can recognize and process them seamlessly. To augment the search precision, a set of filters has been incorporated. Users can customize their search based on predicted pathogenicity scores or narrow down the results using specific labels like ‘likely benign’, ‘likely pathogenic’, or ‘ambiguous’.

Transitioning to the results page, users are greeted with tables, one for each matched protein. These tables, apart from being informative, are interactive. Each table header is clickable, giving users the autonomy to show and hide the contents of individual tables on-the-fly. The data within them provides comprehensive details on variants, showcasing them in a HGVS format and supplementing them with the respective AlphaMissense score and classification. A ‘Download’ option is available for those inclined towards a deeper analysis or external processing of the retrieved data. Clicking on instantly prepares and presents a zipped file named “AlphaMissenseSearch.zip” to the user. Inside, a tab-separated values file contains a set of information columns ranging from UniProt ACC to pathogenicity class, ready to be ingested by software like Excel.

The Hotspot page is where AlphaMissence meets structural biology (Fig. 3). By default, the structure and pathogenicity table for CFTR are displayed, but users have the freedom to input another human protein using one of the supported identifier types. Here, the PDBe-Mol* viewer (39), based on the Mol* framework (40), presents the AlphaFold-predicted structure of the protein in exquisite detail. The displayed structure is fully interactive; users can rotate, highlight, and click on various elements. Additionally, pathogenicity scores are embedded within the occupancy column of the PDB file, so it is easier to ascertain the pathogenic potential of specific residues (the displayed occupancy values by the viewer are AlphaMissence scores). This table and structure can also be downloaded for analysis and structural visualization on desktops. We provide a PyMOL plugin for coloring the downloaded structures based on the B-factor column.

**Figure 3:**
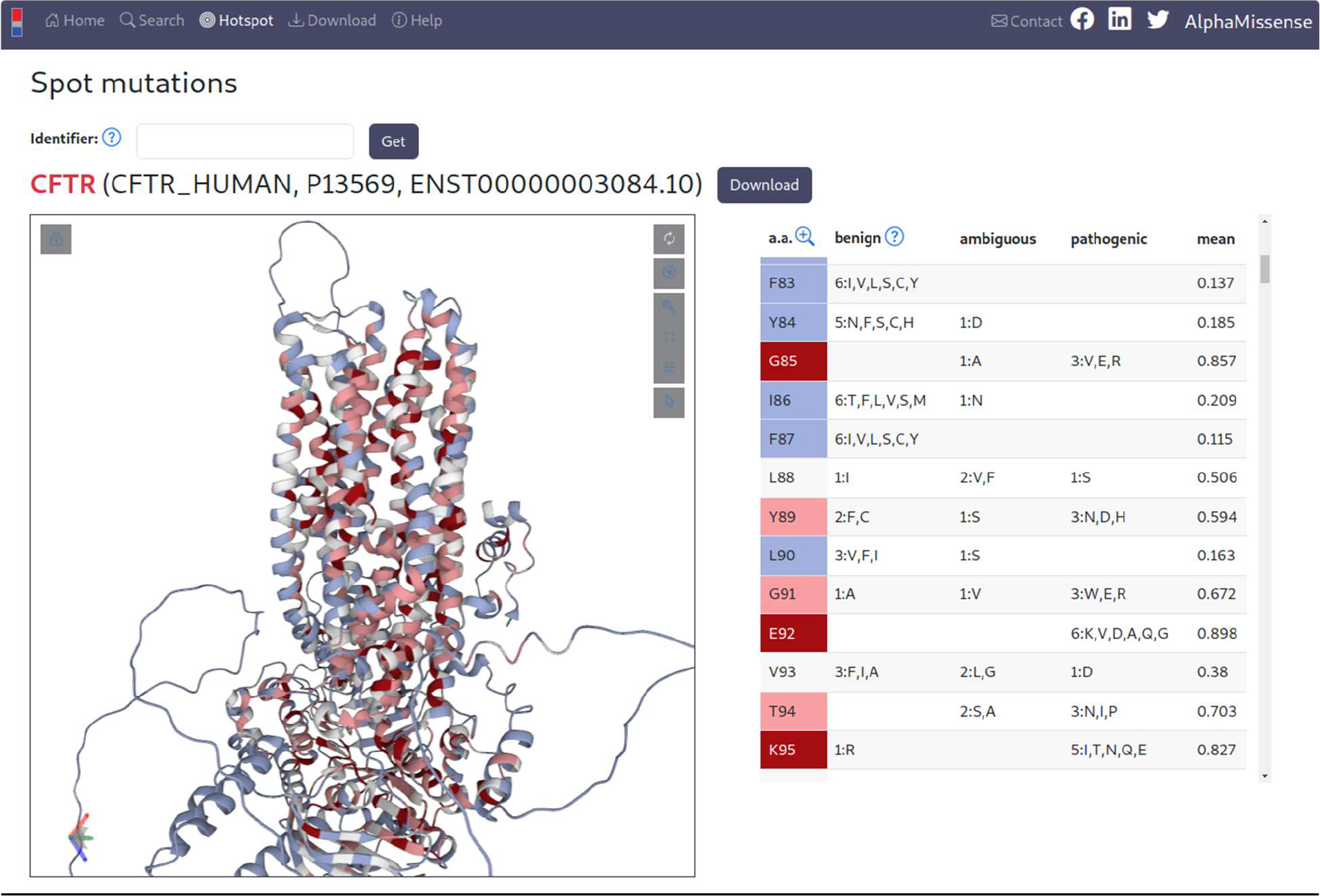
The view of the Hotspot page. Left: PDBe-mol* structure viewer, right: table listing mutations and mean AlphaMissense scores for each residue.

## Conclusion

We embarked on an in-depth analysis of AlphaMissense predictions, ranging from broad protein groups down to the individual CFTR protein. Our objective was to gain insights that would aid the interpretation of predictions for specific target proteins. For benchmarking purposes, we turned to ClinVar, given its substantial repository of curated and reviewed entries. Remarkably, AlphaMissense exhibited consistent performance across various protein categories, evidenced by an MCC value exceeding 0.6 (Table 1). Exceptions arose in scenarios where either the volume of benchmark data was sparse or when the quality of the data was lower. These cases included MemMoRFs and ClinVar’s CFTR data. Our results indicate AlphaMissense performing well when comparing to the CFTR2 database that is in contrast with the study of McDonald *et al*. (41). The differences likely arise from our exclusion of CFTR2 entries with unknown consequences and ambiguous AlphaMissense predictions. We also emphasize that while AlphaFold’s pLDDT scores can provide insights into AlphaMissense performance for specific proteins or protein groups, assessing the rationality of their structures might further indicate the reliability of lower AlphaMissense predictions (lowAF in Table 1).

Both within ClinVar and the AlphaMissense predictions, benign mutations typically outnumbered their pathogenic counterparts by a factor of approximately two, in several protein groups. Intriguing deviations from this trend were noted in groups such as mitochondrial proteins, housekeeping genes, transmembrane regions of membrane proteins, and IBS residues that pattern aligns with expectations. The IBS dataset, with its notably high pathogenic frequency, exclusively contains functional positions (Table 1). The pathogenicity of CFTR coupling helices were also predicted with remarkable congruency with CFTR2 data (Fig. 2e,f). These observations accentuate the potential of AlphaMissense predictions as a valuable tool for aiding the identification of functionally crucial sites. To facilitate hotspot detection and access to AlphaMissense data, we established a dedicated web resource available at https://alphamissense.hegelab.org, which also provides structure files with mapped AlphaMissense scores for visualization, e.g. in PyMOL with our coloring plugin *coloram*.*py*, for facilitating local analysis. These enhancements crucially aid in mutational hotspot detection, paving the way for more detailed and user-friendly analyses.

## Acknowledgments

This work has been supported by the National Research, Development and Innovation Office (grant number: K 137610). Thanks to Mihaly Varadi and Adam Midlik (EMBL EBI) for their help with PDBe-Mol*.

